# GRIN2B disease-associated mutations disrupt the function of BK channels and NMDA receptor signalling nanodomains

**DOI:** 10.1101/2024.10.02.616357

**Authors:** Rebeca Martínez-Lázaro, Teresa Minguez-Viñas, Diego Alvarez de la Rosa, David Bartolomé-Martín, Teresa Giraldez

## Abstract

Large conductance calcium-activated potassium channels (BK channels) are unique in their ability to respond to two distinct physiological stimuli: intracellular Ca^2+^ and membrane depolarization. In neurons, these channels are activated through a coordinated response to both signals; however, for BK channels to respond to physiological voltage changes, elevated concentrations of intracellular Ca^2+^ (ranging from 1 to 10 μM) are necessary. As a result, it is believed that BK channels are typically localized within nanodomains near Ca^2+^ sources (approximately 20-50 nm), such as N-methyl-D-aspartate receptors (NMDARs). Since the first evidence of NMDAR-BK channel coupling reported by Isaacson and Murphy in 2001 in the olfactory bulb, further studies have identified functional coupling between NMDARs and BK channels in other regions of the brain, emphasizing their importance in neuronal function. Mutations in the genes encoding NMDAR subunits have been directly linked to various neurodevelopmental disorders, including intellectual disability, epilepsy, and autism spectrum disorders. For instance, mutations such as V15M and V618G in the GRIN2B gene, which encodes the GluN2B subunit of NMDARs, are implicated in the pathogenesis of early infantile epileptic encephalopathy (EIEE27). Here, we explored the effects of these two GluN2B mutations on NMDAR-BK channel coupling, employing a combination of electrophysiological, biochemical, and imaging techniques. Taken together, our results demonstrate that mutation V618G specifically disrupts NMDAR-BK complex formation, impairing functional coupling, in spite of robust individual channel expression in the membrane. These results provide a potential mechanistic basis for EIEE27 pathophysiology and uncover new clues about NMDAR-BK complex formation.

## Introduction

Large conductance Ca^2+^- and voltage-activated K^+^ channels (KCa1.1, BK, MaxiK or slo1) are expressed in cellular membranes as homotetramers of α subunits encoded by the KCNMA1 (Slo1) gene (Cui et al., 2009; Latorre et al., 2017). BK channels have a wide range of specialised physiological functions across various excitable and non-excitable tissues, such as muscle, kidney, gastrointestinal tract, salivary glands, and bone (Echeverria et al., 2024). In the central nervous system (CNS), they are predominantly found in the soma, axons, and synaptic terminals of cells from regions including the olfactory system, neocortex, basal ganglia, hippocampus, and thalamus (Kshatri et al., 2018; Trimmer, 2015). Within excitable cells, BK channels are primarily involved in shaping the action potential and regulating firing frequency, as well as neurotransmitter release (Bean, 2007; Contet et al., 2016; Storm, 1987). In addition, they have been shown to regulate synaptic transmission and plasticity (Gomez et al., 2021; Zhang et al., 2018).

A key physiological characteristic of BK channel is the dependence of simultaneous membrane depolarization and an increase in intracellular Ca^2+^ levels for activation (Marty, 1981; Pallotta et al., 1981). In many cell types, BK channel activation relies on localized Ca^2+^ rises that reach micromolar concentrations, significantly higher than the typical resting cytosolic levels of 100 to 300 nM (Fakler & Adelman, 2008). Physiologically, BK channels are often situated near other proteins that serve as intracellular Ca^2+^ sources, functioning within specialized Ca^2+^ nano- or micro-domains (Gonzalez-Hernandez et al., 2023; Shah et al., 2021). In synapses, the association of BK to N-methyl-D-aspartate receptors (NMDAR) into Ca^2+^ nanodomains has been proposed to be involved in regulation of synaptic transmission and plasticity (Gomez et al., 2021; Isaacson & Murphy, 2001; Tazerart et al., 2022; Zhang et al., 2018). NMDAR are heterotetrameric ligand-gated ion channels that belong to the family of ionotropic glutamate receptors (iGluR), together with the α-amino-3-hydroxy-5-methyl-4-isoxazolepropionic acid (AMPAR) and kainate receptors (Hansen et al., 2021; Reiner & Levitz, 2018). In physiological conditions these receptors mediate the inflow of Na^+^ and Ca^2+^ and outflow K^+^ (Hansen et al., 2018). NMDARs facilitate the regulated and gradual influx of Ca^2+^ into cells in response to neuronal stimuli, making them essential for processes like synaptic plasticity, learning, memory, and other advanced cognitive functions (Paoletti et al., 2013). Their physiological importance is highlighted by the association between NMDAR dysfunction and a range of neurological and psychiatric disorders, such as Alzheimer’s (Mota et al., 2014) and Huntington (Fernandes & Raymond, 2009) disease, schizophrenia and stroke (Paoletti et al., 2013), as well as major depressive disorder (Molero et al., 2018).

NMDARs assemble at the neuronal membrane as tetramers of various subunit combinations. So far, seven homologous NMDAR subunits have been identified: the essential GluN1/NR1 subunit, four GluN2/NR2 subunits (GluN2A, GluN2B, GluN2C, and GluN2D), and two GluN3/NR3 subunits (GluN3A and GluN3B). The GluN1 and GluN3 subunits have binding sites for co-agonists, such as glycine or D-serine, while the GluN2 subunits feature a binding site for the agonist glutamate (Paoletti et al., 2013). NMDARs configurations include two GluN1 and two GluN2 subunits (GluN1/GluN2), di-heteromeric GluN1/GluN3, or tri-heteromeric GluN1/GluN2/GluN3, contributing to the wide functional diversity of NMDARs in the CNS (Hansen et al., 2018). Additionally, the expression of these genes is regulated both spatially and temporally, further enhancing NMDAR heterogeneity throughout the brain (Paoletti et al., 2013). NMDARs are highly responsive to glutamate, with a half-maximal effective concentration in the micromolar range, and they undergo voltage-dependent blockade by Mg^2+^ ions (Mayer et al., 1984; Nowak et al., 1984). Their slow gating kinetics (Lester et al., 1990) and notable Ca^2+^ permeability (MacDermott et al., 1986; Mayer & Westbrook, 1987) allow postsynaptic NMDARs to effectively sense and interpret the simultaneous activity of both presynaptic and postsynaptic neurons. Specifically, glutamate released from the presynaptic neuron binds to the receptor, while depolarization of the postsynaptic membrane via AMPARs alleviates the Mg^2+^ block. This coordination activates the NMDARs, enabling Ca^2+^ influx through the channel and initiating signalling cascades that can influence synaptic plasticity (Paoletti et al., 2013). Consequently, any regulatory mechanisms that affect Ca^2+^ entry through NMDARs are likely to alter neuronal plasticity and its associated effects.

The initial discovery that Ca^2+^ influx through NMDARs could activate BK channels in specific NMDAR-BK nanodomains was reported in the olfactory bulb (Isaacson & Murphy, 2001), highlighting how Ca^2+^ from glutamate-activated NMDARs triggers BK outward currents. Subsequent research by Zhang and colleagues indicated that this functional relationship might also occur in other brain regions, as evidenced by the co-immunoprecipitation of BK and NMDAR in the hippocampus, cortex, cerebellum, striatum, and thalamus (Zhang et al., 2018). The functionality of these interactions has been examined using whole-cell patch-clamp recordings following the application of glutamate or NMDA at either the neuronal soma (Zhang et al., 2018) or dendrites (Gomez et al., 2021). Notably, the activation of BK by NMDARs in dendrites has been shown in layer 5 pyramidal neurons from cortex (Gomez et al., 2021; Mitchell et al., 2023; Tazerart et al., 2022), and more recently in the CA3 region of the hippocampus, cerebellum, and amygdala (Reyes-Carrión et al., 2023). NMDAR-BK complexes have been identified at both extrasynaptic and postsynaptic terminals (Gomez et al., 2021; Isaacson & Murphy, 2001; Tazerart et al., 2022; Zhang et al., 2018). Within these nanodomains, Ca^2+^ entry through activated NMDARs opens BK channels, resulting in the hyperpolarization of the adjacent plasma membrane and closure of NMDAR channels by restoring Mg^2+^ block. Because BK channel activation blunts NMDAR-mediated excitatory responses, it provides a negative feedback mechanism that may modulate excitability, synaptic transmission or plasticity, depending on the location of these associations within the neuron (Gomez et al., 2021; Shah et al., 2021). Although the mechanisms underlying the association between BK and NMDAR in the nanodomains remain largely unexplored, it has been shown that the isolated GluN1 cytosolic regions directly interact in vitro with a synthesised peptide of the BKL S0-S1 loop region. In addition, the NMDAR-BK interaction is competitively diminished by a synthesised peptide from BKL S0-S1 loop (Zhang et al., 2018). The role of the GluN2 subunits in regulating the formation or function of NMDAR-BK nanodomains remains unclear.

In recent years, different studies have discovered inherited and de novo mutations in genes encoding NMDAR that are directly related to neurodevelopmental disorders, such as mental retardation, intellectual disability, epilepsy and autism spectrum disorders (Lemke et al., 2014; Swanger et al., 2016). Human mutations on the GRIN2B gene have been linked to two types of autosomal dominant neurodevelopmental disorders: EIEE27 (epileptic encephalopathy, early infantile, 27) and MRD6 (mental retardation, autosomal dominant 6) (Lemke et al., 2014; Swanger et al., 2016). Both syndromes show delayed psychomotor development, intellectual disability, seizures, hypotonia, abnormal movements, and autistic features. Moreover, phenotypes of both syndromes are highly variable among patients, ranging from mild intellectual disability without seizures to encephalopathy. We hypothesised that, based on the growing evidence that NMDAR-BK associations play a relevant role in many neuronal types, regulating synaptic function, some of these disease-related mutations may result in alterations of NMDAR-BK functional associations. In this work we have focused on studying the effect of mutations, directly linked to EIEE27, on NMDAR-BK associations. Using a combination of electrophysiology and Ca^2+^ imaging, molecular biology and protein biochemistry, total internal reflection microscopy and superresolution microscopy, we now show that two mutations in the GluN2B subunit (V15M and V618G), linked to EIEE27 alter the functional association of NMDAR and BK in nanodomains, using different mechanisms. The V15M mutation does not impact the efficiency of NMDAR-BK coupling, but it significantly reduces the membrane levels of NMDARs. This leads to fewer functional NMDAR-BK complexes and likely alters the molecular ratio of BK to NMDAR within those complexes. In contrast, the V618G mutation specifically affects the efficiency of NMDAR-BK coupling by changing the composition and size of the NMDAR-BK nanodomains, likely through modifications in their molecular interactions. Notably, our data indicate that the formation of NMDAR-BK macrocomplexes may not solely depend on GluN1-BK interactions, as previously suggested (Zhang et al., 2018). Our findings with disease-related mutations indicate that the functionality of NMDAR-BK nanodomains is influenced by cluster size and the molecular ratio of NMDAR to BK channels, rather than the distance between proteins within the nanodomain. This suggests a potential mechanism underlying the functional variability of NMDAR-BK macrocomplexes.

## Methods

### Cell culture, transfection, cDNA constructs and mutagenesis

HEK293T cells (American Type Culture Collection no. CRL-3216) were grown in Dulbecco’s Modified Eagle Medium (DMEM, Sigma-Aldrich) supplemented with 10% foetal bovine serum (FBS, Sigma-Aldrich), 1% penicillin-streptomycin (Thermo-Fisher Scientific) and Mycozap^TM^ (Lonza). Cells used for imagining and electrophysiology experiments were seeded on poly-lysine-treated glass coverslips to promote cell attachment. Cells grown to 60-80% confluency were transfected with the indicated plasmid combinations using jetPRIME® reagent (Polyplus) following the manufacturer’s instructions. Four hours after adding the transfection mix, medium was replaced, and cells were incubated at 37°C for 24-48h. The following plasmids were used for transient cell transfections: pEYFP-NR1a, encoding the NMDAR-NR1 subunit tagged with an enhanced yellow protein (EYFP) (Addgene plasmid #17928, http://n2t.net/addgene:17928; RRID:Addgene_17928; (Luo et al., 2002)); pEGFP-NR2B, encoding the NMDAR-NR2B subunit tagged with an enhanced green fluorescent protein (EGFP) (Addgene plasmid #17925; http://n2t.net/addgene:17925; RRID:Addgene_17925; (Luo et al., 2002)); pEGFP-NR2B mutants, encoding for NR2B mutant subunits tagged with an enhanced green fluorescent protein; pBNJ_hsloTAG, encoding for BKα subunits tagged with a DYKDDDDKD flag (TAG) (Giraldez et al., 2005); pcDNA3-EGFP encoding for an enhanced green fluorescent protein (Addgene plasmid #13031; http://n2t.net/addgene:13031; RRID:Addgene_13031); Lyn-R-GECO1 (gift from Won Do Heo (Addgene plasmid # 120410 ; http://n2t.net/addgene:120410 ; RRID:Addgene_120410; (Kim et al., 2016)). All fluorescently tagged NMDAR plasmids were a gift from Stefano Vicini, Georgetown University School of Medicine, Washington, DC (Gomez et al., 2021; Vicini et al., 1998). The transfection ratio of GluN1:GluN2B was in most of the experiments 1:3. When co-transfected with BK channels, BK:GluN1:GluN2B transfection ratio was 1:1:3, except for superresolution imaging experiments in which the transfection ratio was 1:1:2.

### Proximity Ligation Assay (PLA)

PLA was performed using the DuoLink Kit (Sigma-Aldrich) including Duolink In Situ Detection Reagents Red (#DUO92008, Sigma Aldrich). Additional reagents Duolink In Situ PLA Probe Anti-Rabbit PLUS (#DUO92002, Sigma Aldrich) and Duolink In Situ PLA Probe Anti-Mouse MINUS (#DUO92004, Sigma Aldrich) were used. HEK293T cells expressing different combinations of NMDAR and BK channels were fixed with 4% paraformaldehyde for 20 min, permeabilized, and then blocked for 1 h at 37 °C to avoid nonspecific antibody binding. The BK channel was detected using a rabbit polyclonal anti-MaxiK channel α subunit primary antibody (1:200, no. ab219072; Abcam). GluN1, GluN2A, and GluN2B subunits of NMDAR were detected using goat polyclonal primary antibodies anti-NMDAR1 (1:200, ref. NB100-41105; Novus Biologicals), mouse monoclonal anti-NMDAe1 (1:200, ref. sc-515148; Santa Cruz Biotechnology), and anti-NMDAe2 (1:200, ref. sc-365597; Santa Cruz Biotechnology), respectively. Secondary antibodies conjugated with oligonucleotides were supplied with the PLA DuoLink Kit. Controls consisted of non transfected HEK293T cells or cells expressing individually the BK alpha subunit or single NMDAR subunits. Images were acquired on a Leica SP8 inverted confocal microscope and image analysis was performed using the Duolink Image Tool (Sigma-Aldrich) and Fiji software. The PLA technique allows the detection of protein–protein interactions (less than 40 nm) as quantifiable fluorescent dots (Gomez et al., 2021). About 120 cells were chosen randomly in 10 different fields from four independent experiments. Nuclei are then automatically detected and cytoplasm size is estimated, allowing single cell statistical analysis of PLA signal levels. Sizes in pixels were transformed into square microns and PLA signals were normalised to cell area. Figures were graphed using Prism 10 (GraphPad).

### Electrophysiology

HEK293T cells were grown on 18-mm poly-lysine-treated glass coverslips and transfected as described above using the indicated combinations of plasmids. Macroscopic currents were recorded at room temperature (21-23°C) using the whole-cell patch-clamp technique with an Axopatch-700B patch-clamp amplifier (Molecular Devices, Foster City, CA, USA) as described previously (Gomez et al., 2021). Recording pipettes were pulled from a 1.5 mm outside diameter x 0.86 mm inside diameter x 100 mm length borosilicate capillary tubes (#30-0057, Harvard Apparatus, Cambridge, UK) using a programmable patch micropipette puller (Model P-97 Brown-Flaming, Sutter Instruments Co., USA). Micropipette resistance was 5-8 MΩ when filled with the internal solution (145 mM K-gluconate, 5 mM Mg-ATP, 1 mM EGTA, 0.2 mM Na-GTP, 10 mM HEPES; pH 7.4) and immersed in the extracellular solution (145 mM NaCl, 5 mM HEPES, 10 mM glucose, 5 mM KCl, 2 mM CaCl2, 10 μ glycine; pH 7.4) (Gomez et al., 2021). Electrophysiological recordings were obtained using the setup described above and Clampex software (pClamp suite, Molecular Devices, California, USA) at 10.000 Hz acquisition rate and 5 kHz low pass filter.

### Intracellular Ca^2+^ fluorescence recordings

Cells were imaged using a NIKON Eclipse Ti-U microscope equipped with a Lumencor Spectra X LED, featuring a green 540 nm LED line, a 40x dry objective with a numerical aperture (NA) of 0.65, an ET - mCherry, Texas Red® (Chroma) filter cube, and an iXon Ultra 888 EM-CCD camera (Andor). Fluorescent cells were patched and recorded as described above. Micro-Manager Open Source Microscopy Software was used for fluorescence data acquisition. Fluorescent cell images were captured in 16-bit format at 4 Hz frequency acquisition. Exposure time was 100 ms. The recordings were synchronised with the amplifier via remote control using Digidata TTL-Outputs (Transistor-Transistor Logic), enabling simultaneous recording of current and fluorescence. Electrophysiology data was analysed using pCLAMP 11 software (Molecular Devices), while fluorescence data was processed with ImageJ. Briefly, images were background subtracted with the ImageJ “BG subtraction from ROI” plugin and the “Time Series Analyzer V3” plugin was used to obtain the fluorescence intensity over time. The changes in fluorescence intensity compared to the baseline fluorescence levels before the application of glutamate (delta F/F0) were graphed against time.

### Cell lysis, protein purification and concentration determination

Total protein extracts were obtained from transfected HEK293T cells resuspended in 50 μl of lysis buffer (20 mM Tris pH 8.0, 100 mM NaCl, 1 mM EDTA, 0.05 % Triton X-100) supplemented with protease inhibitors (Roche). After incubating for 5 min on ice, the cell suspensions were centrifuged for 10 min at 14,000 xg at 4°C. Protein concentration was determined using the bicinchoninic acid assay (BCA) (Smith et al., 1985).

### Cell surface biotinylation

Biotinylation and recovery of membrane proteins was carried out essentially as described before (Alvarez de la Rosa et al., 2002). Experiments were carried out at 4°C, to minimise cell detachment from the plates and stop membrane trafficking. Transfected HEK293T cells were first washed with ice-cold DMEM and then twice with phosphate-buffered saline (PBS) containing 0.1 mM CaClL and 1.0 mM MgClL (PBS-Mg-Ca solution). Cells were then incubated with EZ Link Sulfo-NHS-SS-biotin (ThermoFischer) at a working concentration of 1.5 mg/ml, freshly diluted into the biotinylation buffer (10 mM triethanolamine at pH 7.5, with 2 mM CaClL and 150 mM NaCl). This incubation was performed twice for 25 minutes at 4°C with very gentle horizontal motion to ensure thorough mixing. Cells were then rinsed twice with PBS-Ca-Mg containing 100 mM glycine and then washed in this buffer for 20 minutes at 4°C to quench all unreacted biotin. Cells monolayers were then rinsed twice more with PBS-Ca-Mg and proteins solubilized in 1 ml of lysis buffer (1.0% Triton X-100, 150 mM NaCl, 5 mM EDTA, and 50 mM Tris pH 7.5) on ice for 60 minutes. Cells were then scraped and lysates clarified by centrifugation at 14,000 xg for 10 minutes at 4°C. Following this, 50-100 μ of packed streptavidin-agarose beads were added to each 900 μ of supernatant and incubated overnight at 4°C with end-over-end rotation. The beads were then washed three times with lysis buffer, twice with high-salt wash buffer (similar to the lysis buffer but contains 0.1% Triton X-100 and 500 mM NaCl), and once with no-salt wash buffer (10 mM Tris, pH 7.5). Proteins were eluted from the beads in 50-100 μ of sodium dodecyl sulfate (SDS)-containing sample buffer.

### SDS-PAGE and Western Blot

Protein samples were resolved by SDS-PAGE. Samples (1 μg/μl) were prepared by mixing protein extracts with 6X Laemmli Buffer (4% SDS, 20% glycerol, 10% 2-mercaptoethanol, 0.004% bromophenol blue and 0.125 M Tris HCl, pH 6.8). SDS-PAGE was performed in Mini-PROTEAN® TGX Stain-Free™ Precast gels (Bio-Rad). The stain-free system allowed *in situ* protein photoactivation after electrophoresis for total protein load visualisation, quantification and normalisation. Running buffer was prepared by dilution of a 10X stock (25 mM Tris, 192 mM glycine, 0.1 % SDS) in MilliQ H_2_O. Electrophoresis was carried out at a constant voltage of 150 V for approximately 1h. Proteins were transferred to a polyvinylidene difluoride (PVDF) membrane using a Trans-Blot^®^ Turbo™ Transfer Starter System for Western Blot analysis at 1.3 A and 25 V for 10 mins. Proteins of interest were visualised and detected in the PVDF membranes employing the primary antibodies mouse anti-GluN2B (#365597, Santa Cruz Biotechnology; 1:1000 dilution), mouse anti-α1 Na^+^,K^+^-ATPase (ATP1A1) monoclonal antibody F-2 (sc-514614, Santa Cruz Biotechnology; 1:1000 dilution) mouse anti-tubulin monoclonal antibody AA13 (T8203, Sigma Aldrich; 1 μ ml), followed by a secondary anti-mouse horseradish peroxidase-conjugated antibody made in goat (P0447, Dako; 1:20,000 dilution). Chemiluminescence signals were recorded in a Chemidoc imaging system (Bio-Rad) and quantified using Image Lab software 6.0 (Bio-Rad).

### Total internal Reflection Microscopy (TIRFM)

Total internal reflection fluorescence microscopy (TIRFM) is an optical technique that enables the excitation of fluorophores within a very thin axial region (200 nm) close to the coverslip, known as the ‘optical section’. TIRFM was performed in a motorised Nikon Eclipse Ti microscope equipped with a 100x immersion objective. The setup included a laser unit with a DPSS 488 laser and a 647 nm fiber laser. GFP-tagged GluN2B was visualised using 1% DPSS 488 laser. Images were captured using an Orca Flash 4.0 CMOS camera (Hamamatsu, Shizuoka, Japan). To quantify the degree of GluN2B-EGFP expression at plasma membrane, TIRF images were background subtracted using ImageJ and normalised to their integrated density and exposition time (ranging 5-100 ms).

### Direct Stochastic Optical Reconstruction Microscopy (STORM)

STORM imaging was performed on a Nikon N-STORM superresolution system with a Nikon Eclipse Ti inverted microscope equipped with an HP Apo TIRF 100X oil NA 1.49 objective (Nikon), a Perfect Focus System (Nikon), and an ORCA-Flash4.0 V2 Digital CMOS camera C11440 (Hamamatsu). Fluorescence emission was filtered with a 405/488/561/640-nm Laser Quad Band filter cube (TRF89901; Chroma). The imaging buffer specific for STORM microscopy contained 50 mM Tris-HCl (pH 8), 10 mM NaCl, 10% (wt/vol) glucose, 100 mM β mercaptoethylamine, 0.56 mg/ml glucose oxidase, and 34 µg/ml catalase (all reagents from Sigma-Aldrich). Reconstructed images were generated from 5 x 10^4^ acquired frames (2.5 x 10^4^ per channel) using the NIS-Elements software (Nikon). We performed at least three independent transfection experiments for each protein combination shown in this study. For every experiment, we determined the location of hundreds of thousands of molecules. Lateral localization accuracy was estimated, as described previously (Kshatri et al., 2020), as 13 ± 4 nm for Alexa Fluor 647 and 16 ± 6 nm for Alexa Fluor 488. Reconstructed images were filtered to remove background. Quantitative analysis of STORM images was performed using nearest-neighbor distance (NND) and cluster analysis using in-house script based on the k-Nearest Neighbor (k-NN) and the density-based spatial clustering of applications with noise (DBSCAN) algorithms, respectively, similar to previously published work (Kshatri et al., 2020). The clustering properties of the samples were quantified by adjusting the density filtering to 20, 40 or 60-nm radius with a count of 10 molecules (Supplementary Fig. 3). Clusters were classified in three categories: “only red fluorophores”, “only green fluorophores” (we refer to these two types as “homoclusters”, formed by just one fluorophore), and “red and green fluorophores” (referred to as “heteroclusters”, composed by more than one fluorophore). Cluster distributions are represented as plots of the percentage of each cluster type normalised to all clusters (all fluorophores).

## Results

We initially focused our investigation on a set of mutations whose locations within the molecular structure of GluN2B are represented in Fig.1. These mutations were linked to the EIEE27 disorder and reported in the literature (Lemke et al., 2014; Swanger et al., 2016). We selected mutations V618G and V15M for our study, located in the TMD and NTD (Fig. 1). To investigate whether these disease-linked mutations on the GluN2B subunit alter the coupling of NMDAR and BK channels, we performed whole-cell voltage-clamp recordings from HEK293T cells transiently transfected with NMDARs containing the GluN1a subunit together with wild-type or mutant GluN2B subunits, which were co-transfected with or without the BK channel α subunit (Fig. 2).

**Figure 1.**
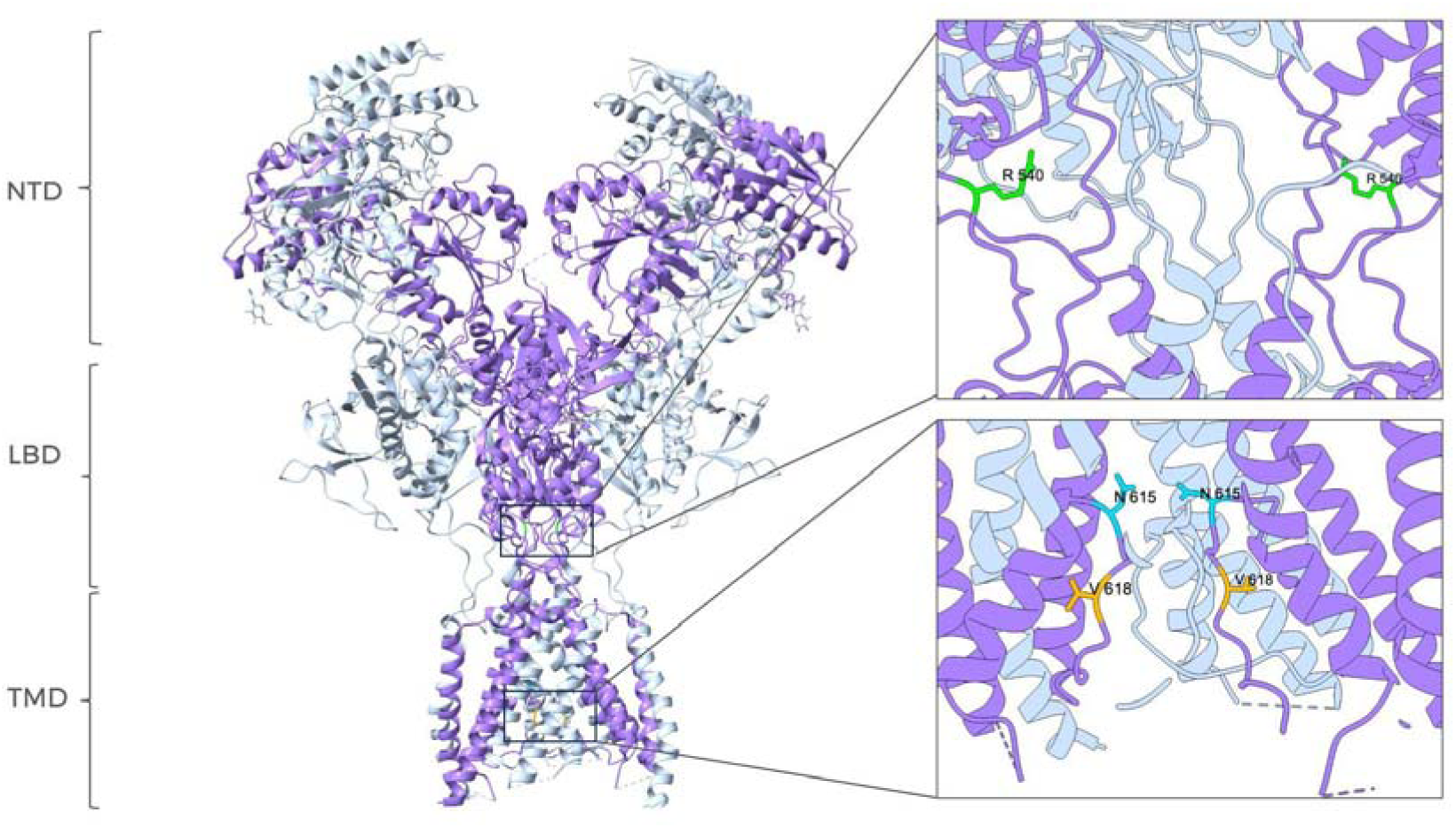
Site location of EIEE27-linked GRIN2B mutations within the structure of NMDAR. (PDB: 7SAA) (Chou et al., 2022). The GluN1 subunit is coloured in grey, GluN2B subunit is depicted in purple. All subunits contain three main regions, indicated on the left side of the figure: N-terminal domain (NTD), ligand-binding domain (LBD) and transmembrane domain (TMD). The position of residue V15 has not been well resolved in the published structure and is not shown. Mutation R540H targets the LBD whereas mutations N615I and V618G are located in the pore lining region within the TMD. The inset shows amplified images of specific regions containing the EIEE27-linked mutations.

**Figure 2.**
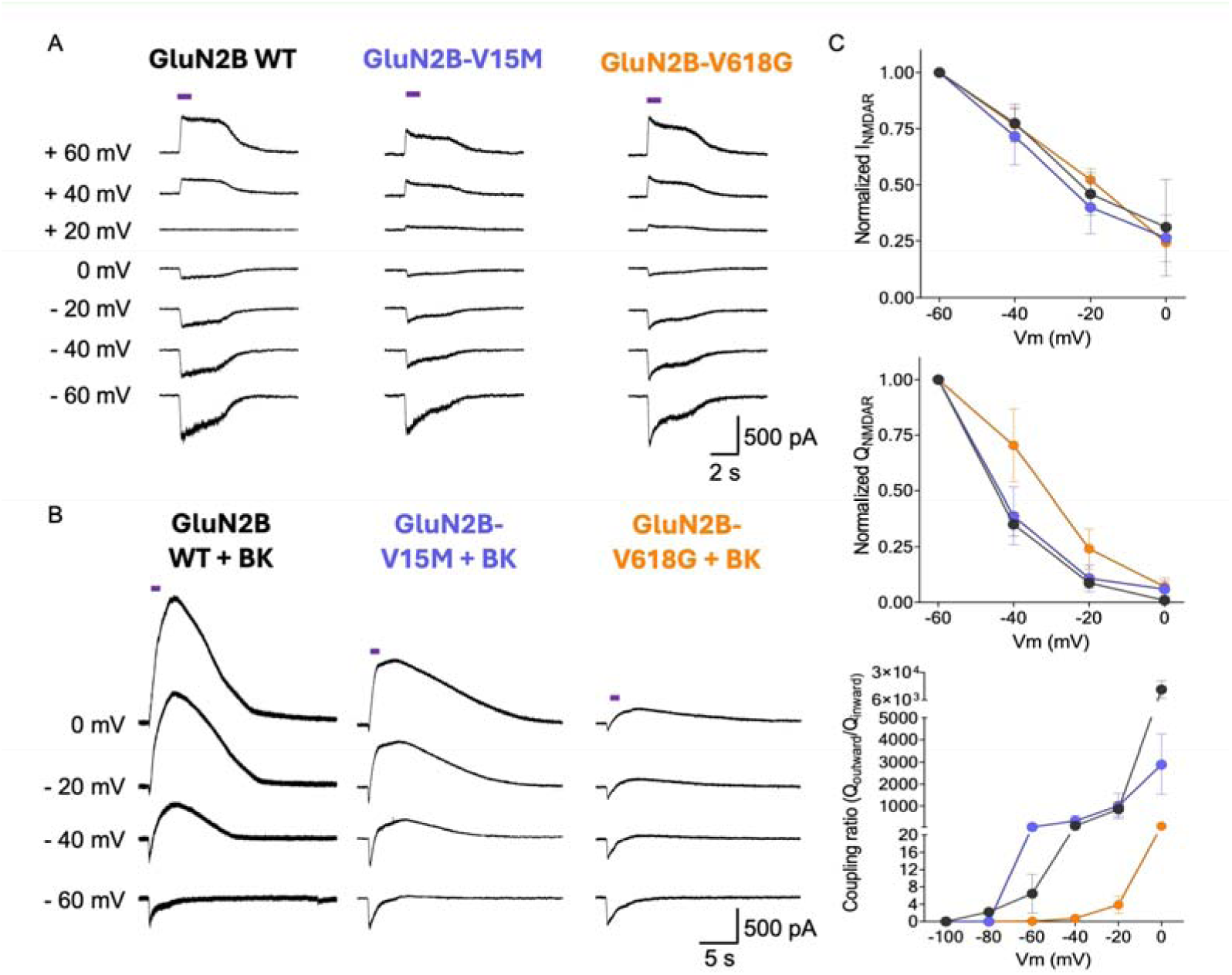
Mutation V618G selectively disrupts functional NMDAR-BK coupling. (A) Representative whole-cell current traces recorded from cells expressing NMDAR combinations GluN1-GluN2B^WT^, GluN1-GluN2B^V15M^, and GluN1-GluN2B^V618G^ alone; and (B) co-expressed with BK after 1 s application of 1mM glutamate (purple square over the traces) evoked at the indicated membrane potentials. (C) Normalised current-voltage (I-V, up) and charge-voltage (Q-V, middle) relationships for NMDAR inward currents from experiments shown in B. The bottom graph represents the efficiency of functional coupling estimated as Qoutward/Qinward relationships vs. voltage for all conditions tested in B. Data points in all graphs represent mean ± SEM; n=5-7.

Cells expressing GluN1-GluN2B^V15M^ or GluN1-GluN2B^V618G^ NMDARs produced inward currents after the application of 1mM glutamate, very similar to those produced by GluN1-GluN2B^WT^ at all potentials studied (Fig. 2A). The characteristics of GluN1-GluN2B^V618G^ were comparable to those reported for these receptors in equivalent experimental conditions (Fedele et al., 2018), whereas to our knowledge the GluN1-GluN2B^V15M^ recordings shown here are the first reported to date.

Voltage-clamp recordings from cells co-expressing GluN1-GluN2B^WT^ receptors with BK channels showed inward currents followed by a slower outward current at holding potentials more positive than −40 mV (Fig. 2B), similar to previously reported NMDAR-activated BK currents (Gomez et al., 2021; Isaacson & Murphy, 2001; Zhang et al., 2018), with clear dependence on membrane voltage (Fig. 2C, bottom). Interestingly, co-expression of GluN1-GluN2B^V15M^ with BK produced comparable results (Fig. 2B, middle panel). However, GluN1-GluN2B^V618G^ failed to activate BK channels as efficiently as GluN1-GluN2B^WT^ or GluN1-GluN2B^V15M^. This finding was consistent with the observed reduction in the net inward current flow produced by the activation of the outward current, which was similar in cells co-expressing BK and GluN1-GluN2B^WT^ as well as GluN1-GluN2B^V15M^ (Fig. 2C, middle graph). In the case of cells co-expressing BK with GluN1-GluN2B^V618G^, this reduction was significantly smaller, resulting in a lower decrease of charge transfer (Gomez et al., 2021). We also quantified the efficacy of NMDAR-to-BK coupling by measuring the ratio between the inward charge and the outward charge, which we refer to as the “coupling ratio”. As shown in Fig. 2C (bottom), recordings from cells co-expressing BK with GluN1-GluN2B^WT^ or with GluN1-GluN2B^V15M^ showed comparable coupling ratios, whereas the values corresponding to cells co-expressing BK with GluN1-GluN2B^V618G^ were significantly smaller. Altogether, these results indicate that mutation V618G in the GluN2B subunit produces selective uncoupling of NMDAR activity from BK when both proteins are coexpressed.

Several studies on the mutation V618G on GluN2B/GRIN2B have been reported (Fedele et al., 2018; Lemke et al., 2014; Vyklicky et al., 2018), although the functional implications of this mutation are not yet fully understood. Valine 618 is a critical and highly conserved residue (Supplementary Fig. 1) located in the linker between the M2 and M3 transmembrane domains, both of which form part of the channel pore lining (Chou et al., 2022). The environment surrounding V618 is highly hydrophobic (Supplementary Fig. 2). Previous reports have shown that this mutation does not alter the receptor’s response to glutamate or glycine, but is associated with reduced NMDAR desensitisation rates, decreased open probability and lower single-channel amplitude (Vyklicky et al., 2018). These kinetic effects are however contrasted by a reduction in Mg^2+^ block (Vyklicky et al., 2018), which has been related to a role of V618 in Mg^2+^ coordination (Fedele et al., 2018). In addition, molecular dynamics studies have proposed that the V618G mutation produces a significant reorientation of the backbone carbonyl groups within the ion filter (Vyklicky et al., 2018), with a significant alteration of the hydrophobicity profile (Supplementary Fig. 2).

Altogether, these observations suggest that the mutation would possibly result in lower selectivity and efficiency of ion transport. This prediction seems however contradicted by two-electrode voltage-clamp experiments in Xenopus oocytes expressing GluN1-GluN2B^V618G^ receptors, which showed increased Ca^2+^ permeability in Mg^2+^-free, NMDG-Cl solutions (Lemke et al., 2014). In contrast, a more recent study shows comparable levels of Ca^2+^ permeation of GluN1-GluN2B^V618G^ compared to wild-type NMDARs (Fedele et al., 2018).

Taking into account the antecedents above mentioned, we reasoned that an alteration in Ca^2+^ permeability could explain the disruption of NMDAR-BK coupling in the macrocomplexes containing GluN1-GluN2B^V618G^ receptors, since the lower availability of Ca^2+^ may activate fewer BK channels in the nanodomain. To assess this possibility, we co-expressed the different NMDAR-BK combinations with Lyn-R-GECO1, a low-affinity red fluorescent genetically-encoded Ca^2+^ indicator for optical imaging fused to a myristoylation signal peptide that targets it to the plasma membrane (Kim et al., 2016). When co-expressed with BK and GluN1-GluN2B^WT^, Lyn-R-GECO1 reported the highest fluorescence and thus, Ca^2+^ permeation, at the most negative potentials recorded (−60 mV), which is consistent with the driving force for Ca^2+^ in our experimental design (0 mM Ca^2+^ intracellular solution vs. 2 mM CaCl_2_ in the extracellular solution). Strikingly, both GluN1-GluN2B^V15M^ and GluN1-GluN2B^V618G^ allowed the entrance of Ca^2+^ to similar extents as GluN1-GluN2B^WT^, as reported by the Lyn-R-GECO1 fluorescence recordings (Fig. 3). This finding suggests that the defective coupling of GluN1-GluN2B^V618G^ with BK channels does not appear to be due to alterations in Ca^2+^ permeability of the mutant receptors. Based on previous findings reporting altered Mg^2+^ permeability in GluN1- GluN2B^V618G^ (Fedele et al., 2018), we confirmed that the 5 mM Mg-ATP included in our solutions was not affecting our results. The coupling ratio obtained in symmetrical Mg^2+^ solutions (5 mM MgCl_2_) was comparable to our initial measurements (data not shown), eliminating the potential impact of abnormal Mg^2+^ permeability on our findings.

**Figure 3.**
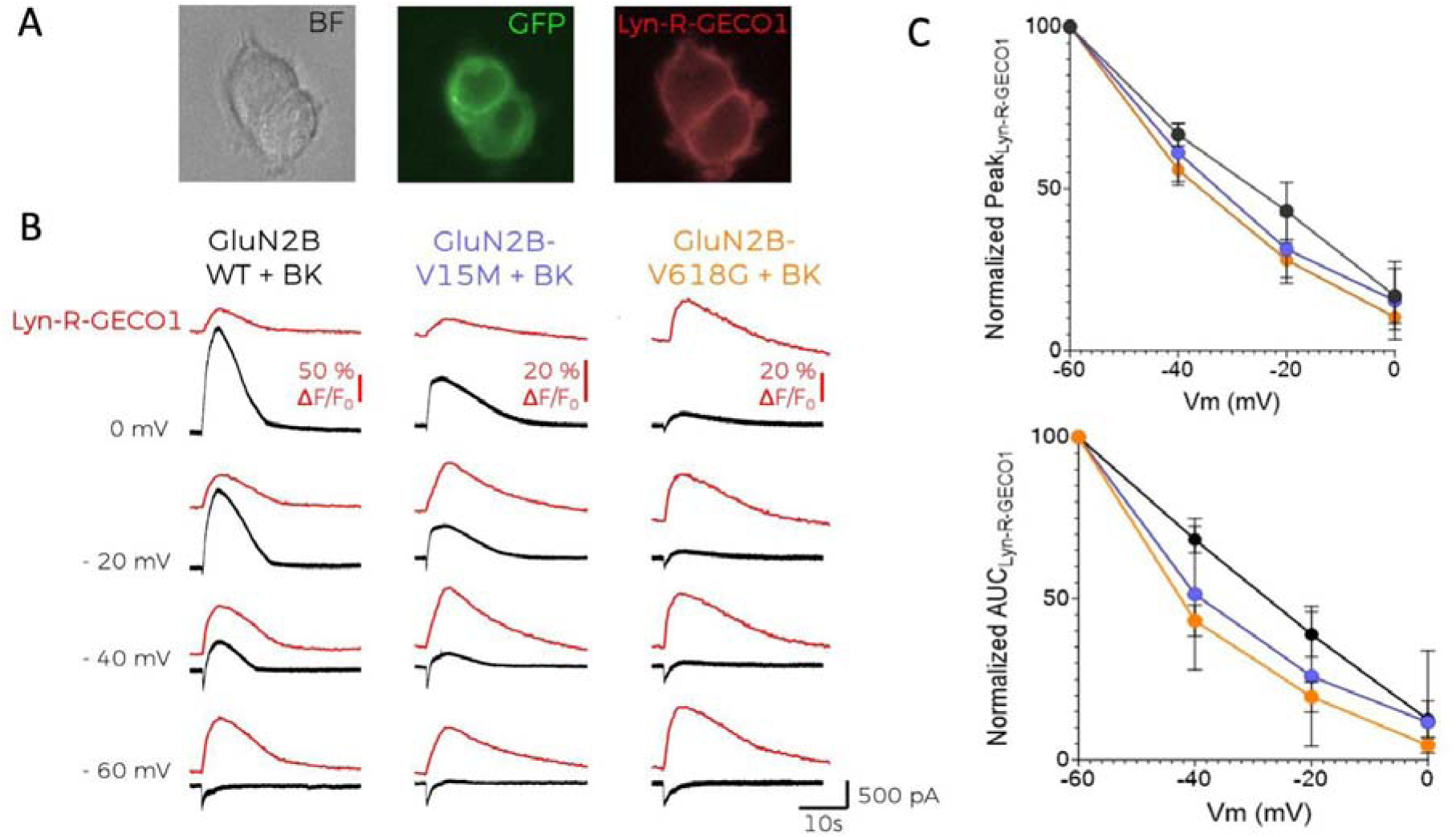
GluN2B variants are permeable to Ca^2+^. (A) Representative microscopy images showing HEK cells co-transfected with BK plus NMDARs containing the GFP-tagged GluN2B variants used in this study, as well as the red fluorescent, membrane-linked Lyn-R-GECO1 Ca^2+^ indicator. Images were obtained in bright field (BF, left panel), with 488 nm excitation light (middle) and with 540 nm excitation light (right panel). (B) Simultaneous whole-cell current (black traces) and normalised fluorescence recordings (ΔF/F_0_, red traces) from cells co-expressing BK channels, the membrane calcium sensor Lyn-R-GECO1 and GluN1-GluN2B^WT^ (left), GluN1-GluN2B^V15M^ (middle) or GluN1-GluN2B^V618G^ (right). Recordings were obtained at the indicated holding potentials after application of 1mM glutamate for 1 s. (C) Graphs represent the averaged maximal peak from Lyn-R-GECO1 recordings (top) or the normalised area under the curve (AUC) (bottom) vs. voltage relationships corresponding to the experiments shown in B, in the absence of BK; Data points represent mean ± SEM; n = 5-7.

Since Ca^2+^ permeability remained unaltered in NMDAR mutants, we reasoned that differences in protein abundance and/or membrane expression of NMDAR subunits could account for the impaired coupling between GluN1-GluN2B^V618G^ and BK channels. Thus, we decided to study overall expression levels by western blot, using biotinylated membrane fractions to specifically study protein membrane abundance (Fig. 4). Remarkably, the analysis of relative protein abundance revealed that the expression levels of V618G were significantly increased in comparison to GluN2B^WT^. In contrast, analysis of the variant V15M revealed significantly diminished expression levels.

**Figure 4.**
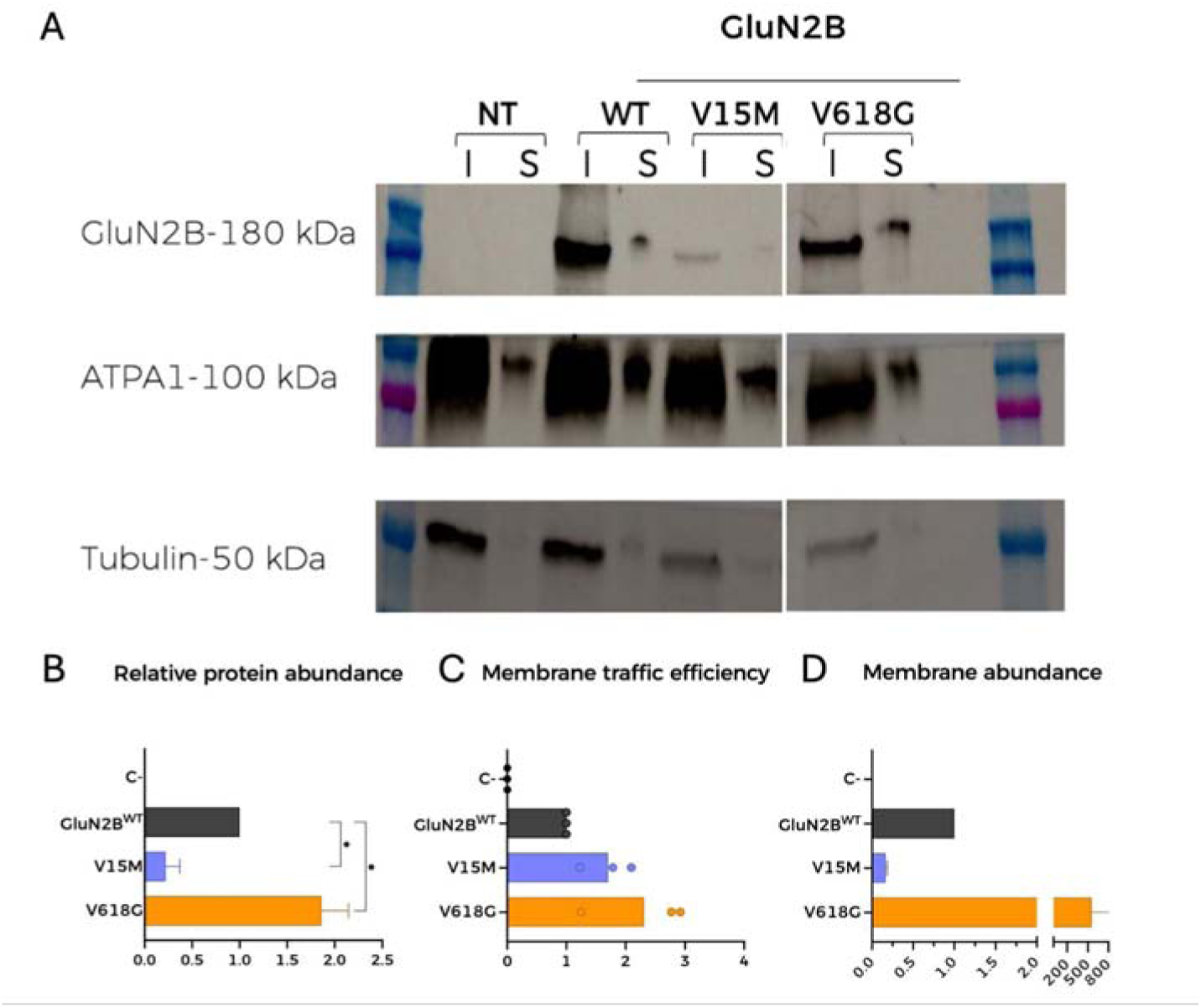
Mutation V618G shows significantly increased membrane abundance without altered trafficking. (A) Representative western blot of NMDAR protein abundance in HEK293T cells transfected with GluN1-GluN2B^WT^, GluN1-GluN2B^V15M^ or GluN1-GluN2B^V618G^. I (input): total lysate abundance; S (surface): biotinylated fractions corresponding to membrane isolates. Total tubulin (lower panel) was used as an internal standard. (B) Left, graph represents the relative protein abundance for NMDAR containing GluN1 and the indicated GluN2B subunits. Data points represent mean ± SEM; n=3, (Dunnet’s multiple comparison test * p> 0.05). Center, graph represents the ratio between GluN2B membrane fractions (S) and total protein expression (I) obtained from experiments in A, for the NMDAR subunit combinations shown. Right, graph representing the total membrane abundance of all NMDAR subunit combinations quantified from biotinylated fractions (S) in A.

Analysis of the GluN2B ratio between the surface fraction retrieved by the beads (S) and the lysate (I) helped understand the efficiency of the traffic to the membrane of GluN1-GluN2B^WT^ and variants (Fig. 4). Notably, GluN1-GluN2B^V15M^ trafficked to the surface membrane very efficiently and so did GluN1-GluN2B^V618G^. This came as a surprise, especially for GluN1-GluN2B^V15M^, which showed the lowest membrane abundance, since the V15 residue falls on the signal peptide. Notably, the membrane expression of GluN1-GluN2B^V618G^ was significantly increased with respect to GluN2B^WT^ (Fig. 4).

The above-mentioned results were independently assessed by using Total Internal Reflection Fluorescence (TIRF) microscopy (Fig. 5). This technique enables the selective excitation of surface-bound fluorophores, allowing to study quantitatively the membrane population of NMDARs containing the different GluN2B variants. HEK293T cells were transfected with GluN1-GluN2B^WT^-EGFP and fluorescent levels were quantified as described (see methods). As shown in Fig. 5B, the GluN1-GluN2B^V618G^ NMDARs showed significantly increased expression at the plasma membrane, whereas GluN1-GluN2B^V15M^ membrane abundance was significantly lower than GluN1- GluN2B^WT^, in agreement with the biotinylation data.

**Figure 5.**
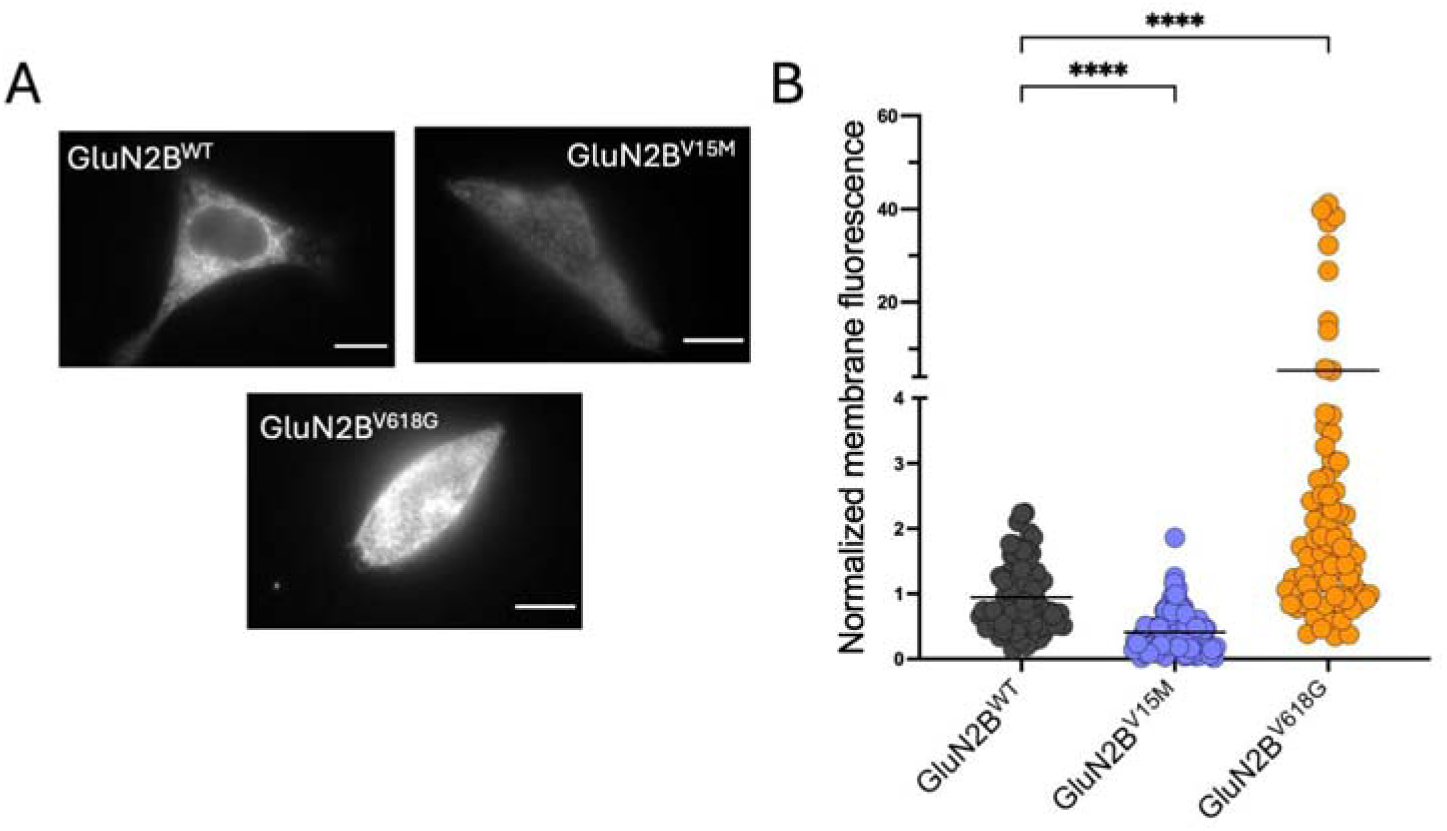
Membrane abundance of different NMDAR combinations measured with TIRF. (A) Representative TIRF microscopy images of HEK293T cells transiently transfected with NMDAR containing combinations of GluN1a and either WT- or mutant-GluN2B subunits. Scale bar=5 μm. (B) Quantitative analysis of TIRFM imaging from cells expressing heteromers of GluN1a and the indicated GluN2B variants, normalised to the fluorescence levels of NMDAR containing WT-GluN2B subunits (black circles). Data represent mean ± SEM (minimum n=30 cell counts per condition and experiment; 3 independent experiments). Statistical analysis was performed with the Kruskal Wallis test (****, p<0.0001).

We then tested whether BK preferentially associates with specific NMDAR subunit variants by using the proximity ligation assay (PLA) (Gomez et al., 2021). This technique is based on the combination of antibody-based protein recognition and nucleotide-based rolling circle amplification, enabling the detection of protein proximity within a radius of 40 nm (Alam, 2018; Gomez et al., 2021). Consistently with our previously published results (Gomez et al. 2021), positive PLA signals were observed for HEK293T cells co-expressing BK and GluN1-GluN2B^WT^, demonstrating that BK channels and NMDARs formed nanodomains in our experimental conditions (Fig. 6A). Cells transfected with GluN1-GluN2B^V15M^ in the presence of BK channels show a marked reduction in the number of positive PLA signals (Fig. 6A-B). This result matched the lower membrane abundance of this variant. Most importantly, PLA signals for cells transfected with GluN1-GluN2B^V618G^ and BK channels were also diminished in comparison with BK and GluN1-GluN2B^WT^. We did not anticipate this result, given the large increase in membrane abundance of GluN1-GluN2B^V618G^ in comparison with GluN1-GluN2B^WT^ (Fig. 4).

**Figure 6.**
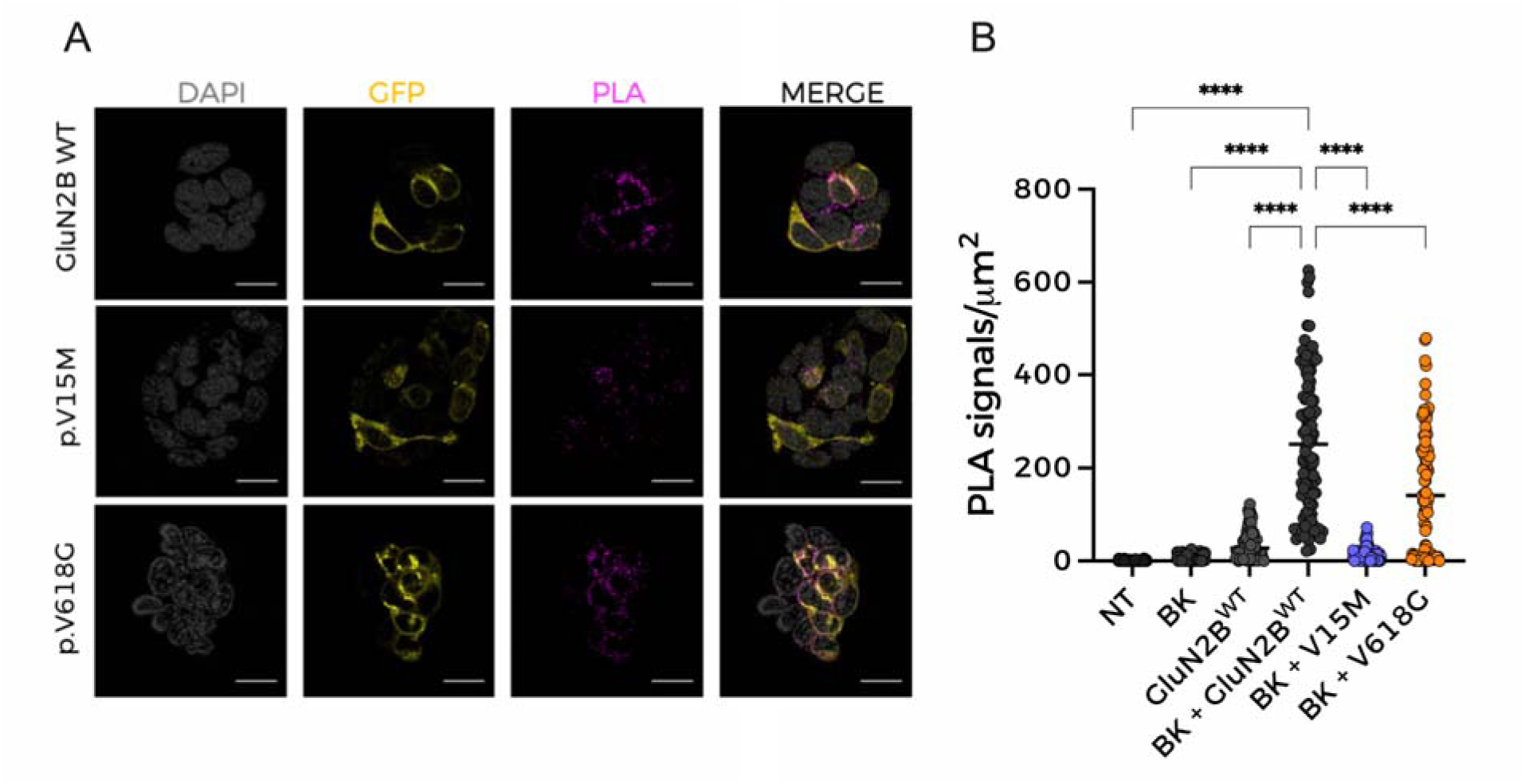
NMDAR containing disease-linked GluN2B subunits show reduced protein-protein complex formation. (A) Representative confocal microscopy images of PLA experiments in HEK293T cells expressing the protein combinations indicated on the left of each row. Each column corresponds to an imaging channel (left, DAPI, 405 nm; middle, NMDAR, 488 nm; right, PLA, 540 nm); merged channels are shown at the far-right column. Scale bar is 20 μm. (B) Quantification of PLA signals/µm^2^ for HEK293T cells transiently expressing BK in the presence of GluN1a combined with the indicated GluN2B subunits: WT, V15M or V618G. Data points represent individual cells, with horizontal bars representing the mean (minimum n=35 cell counts per experiment; 4 independent experiments). Statistical analysis was performed with the Kruskal Wallis test (****, p<0.0001 vs. BK+ GluN1- GluN2B^WT^).

Altogether, our data show that the functional coupling of GluN1-GluN2B^V618G^ receptors to BK channels is significantly diminished as compared with that of GluN1-GluN2B^WT^ or another disease-linked mutant, GluN1-GluN2B^V15M^. This effect occurs despite a higher membrane abundance of GluN1-GluN2B^V618G^, but is consistent with the observed reduction of complex formation between GluN1-GluN2B^V618G^ and BK channels reported by PLA experiments. Two scenarios may possibly explain these results. On one hand, mutant receptors GluN1-GluN2B^V618G^ and BK channels could be located at further distances within the complexes. Another possibility may be any alteration of the multichannel cluster characteristics (size, composition, or a combination of both), when GluN1-GluN2B^V618G^ and BK channels are co-expressed. To discern whether any of these possibilities, or a combination of all of them, may reconcile all our observations and enlighten the cellular mechanism underlying the altered functional coupling of GluN1-GluN2B^V618G^-BK complexes, we used STORM super-resolution microscopy. This technique was combined with TIRF to investigate the spatial organisation of NMDAR-BK complexes at or near the plasma membrane (Fig. 7 and 8) (Kshatri et al., 2020). Close localizations of BK and GluN2B^WT^ were observed, as reflected in the NND distribution analysis, which showed a higher peak at 25–30 nm (Fig. 7B), supporting previous findings indicating that BKα and GluN1-GluN2B^WT^ are in nanoscale proximity (Gomez et al., 2021; Zhang et al., 2018). Surprisingly, the BKα-to-GluN2B distance distribution was practically identical in cells co-expressing BK with GluN1-GluN2B^WT^ or with BK GluN2B^V618G^ (Fig. 7C). These results indicate that the presence of the V618G mutation in the GluN2B subunit is not associated with increased distances between BK and NMDAR in the nanodomains.

**Figure 7.**
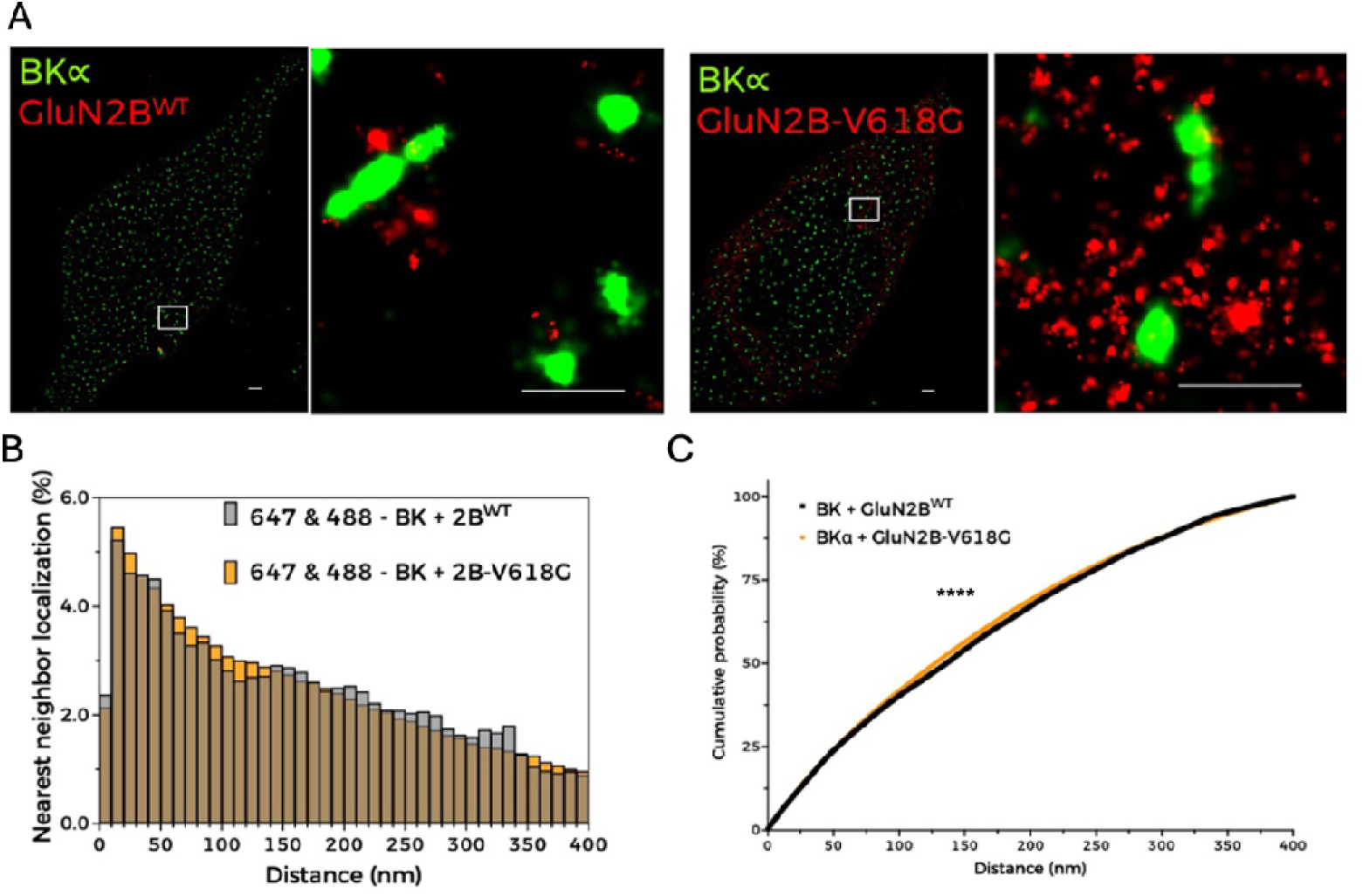
BK and NMDAR are located in nanoscale proximity. (A) Representative STORM images (left; scale bar: 1 µM) and magnified views of areas of interest (right; scale bar: 0.5 µM) showing the spatial distribution of BK_α_ (green, Alexa Fluor 488) and GluN2B (red, Alexa Fluor 647) in HEK293T cells co-expressing BK, GluN1a and either GluN2B^WT^ or GluN2B-V618G. (B) NND analysis from the corresponding dual-label experiments indicated in the graph legend. (C) Cumulative probability analysis of NND distribution (Kolmogorov-Smirnov test ****p<0.0001, D=0.0244).

**Figure 8.**
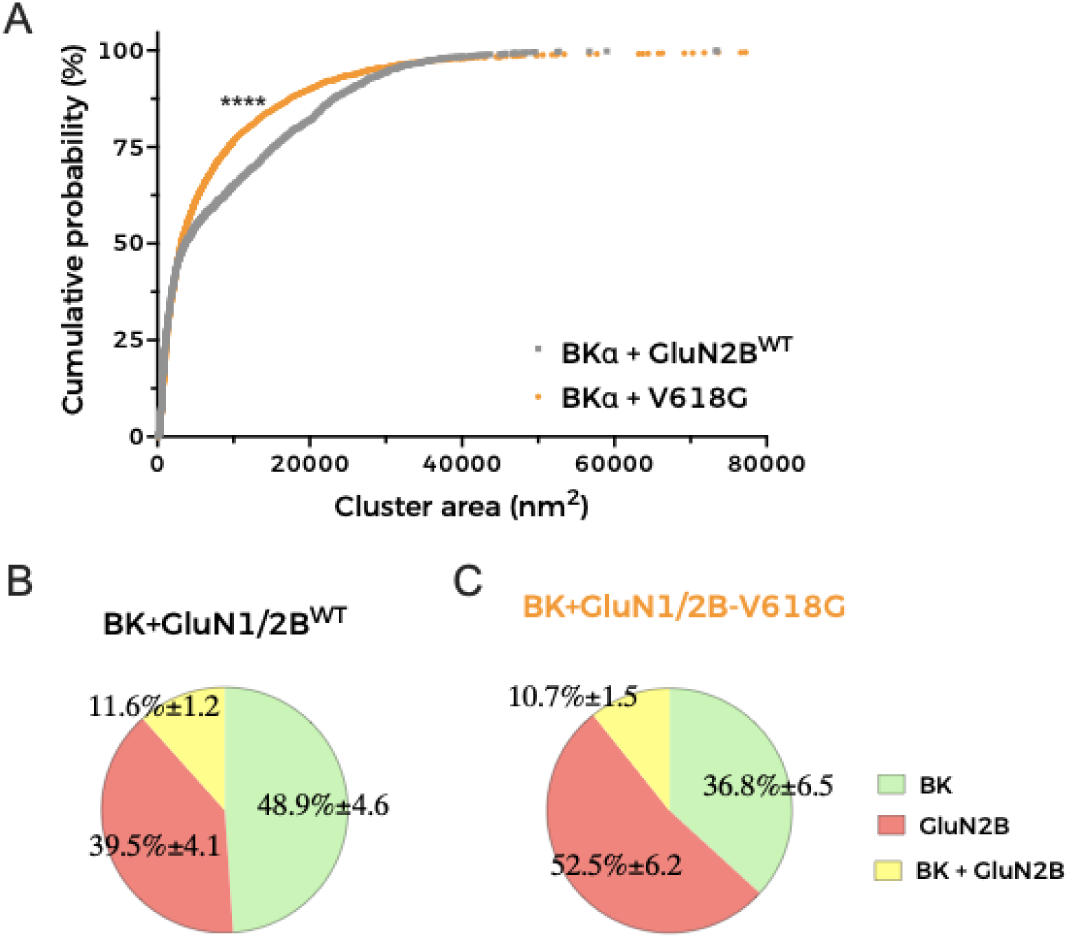
Distribution of clusters in HEK293T cells co-expressing BK and GluN1- GluN2B^WT^ or BK with GluN1-GluN2B^V618G^. Representative cluster analysis data with radius cutoff set to 20 nm (see Supplementary Fig. 3 for a complete description of analyses and data). (A) Cumulative probability analysis of heteroclusters distribution in cells co- expressing BK and either GluN1-GluN2B^WT^ or GluN1-GluN2B^V618G^ (Kolmogorov-Smirnov test ****p<0.0001, D=0.1183). (B-C) Pie charts represent the percentage of clusters with the indicated combinations in cells co-expressing BK and either GluN1-GluN2B^WT^ or GluN1-GluN2B^V618G^. Composition of clusters is colour-coded as described in the graph legend.

Cluster analysis of superresolution data provides useful insights into spatial patterns and associations between proteins (Kshatri et al., 2020; Ricci et al., 2015; Vivas et al., 2017; Zanacchi et al., 2017). We performed this analysis to better understand whether there are any differences in cluster formation between BK/GluN1-GluN2B^WT^ and BK/GluN1-GluN2B^V618G^. We used an in-house software written in Python to identify and calculate areas of clusters with all possible protein combinations in each experimental condition. This analysis applies the DBSCAN algorithm, a data-clustering algorithm that finds core samples of high density and expands clusters from them. This algorithm is based on two parameters set by the experimenter referring to the radius of the core cluster and to the minimum number of particles (Kshatri et al., 2020; Ricci et al., 2015). Based on previous experience, we analysed clusters setting the core size to 10 particles (Kshatri et al., 2020) and generated three full analyses considering radii of 60 nm, 40 nm and 20 nm (Supplementary Fig. 3). A striking result was that, in all conditions tested, BK/GluN1-GluN2B^WT^ heteroclusters were significantly larger than heteroclusters formed by BK and GluN1-GluN2B^V618G^, as inferred from the analysis of the cumulative probability of cluster area (Fig. 8A). In addition, comparison of all the obtained distributions (Supplementary Fig. 3) consistently showed the following: (i) the proportion of NMDAR-BK heteroclusters quantified in cells expressing BK and GluN1-GluN2B^WT^ was very similar to that observed in cells expressing BK and GluN1-GluN2B^V618G^ (Fig. 8B and 8C, yellow); (ii) BK homoclusters (Fig. 8B, red) are more abundant in cells co-expressing BK with GluN1-GluN2B^WT^ than in those co-expressing BK with disease- linked mutant GluN1-GluN2B^V618G^ (Fig. 8B and 8C, red); and (iii) NMDAR homoclusters are more abundant in cells co-expressing BK with GluN1-GluN2B^V618G^ than in those co-expressing BK with GluN1-GluN2B^WT^ (Fig. 8B and 8C, green). The latter observation is consistent with the biotinylation and TIRF data shown in this study (Fig. 4 and Fig. 5). Altogether, these observations lead us to conclude that the proportion of GluN1-GluN2B and BK particles must be different in nanodomains containing BK/GluN1-GluN2B^V618G^ and those containing BK/GluN1-GluN2B^WT^.

## Discussion

In this work we provide additional evidence that BK and NMDAR nanodomains can be functionally reconstituted in a heterologous expression system such as HEK293T, offering a valuable model system to understand the mechanisms underlying formation and function of these channelosomes. Functional coupling of GluN1-GluN2B^WT^ to BK was recorded electrophysiologically and recapitulated the biophysical properties previously described in neurons (Gomez et al., 2021; Isaacson & Murphy, 2001; Zhang et al., 2018). PLA and superresolution analysis showed that BK and GluN1-GluN2B^WT^ are located in nanoscale proximity, showing a sharp NND maximal peak at around 25 nm. The nanoscale proximity of BK channels and NMDARs is a critical aspect of their functional relationship in neurons, facilitating efficient Ca^2+^ signalling and modulating neuronal excitability, synaptic transmission and plasticity (Gomez et al., 2021).

We discovered that two disease-linked mutations in the GluN2B subunit, V15M and V618G, are related to alterations in BK-NMDAR coupling. A striking observation of this study is that NMDARs containing GluN2B^V618G^ subunits showed disrupted functional coupling to BK channel function, as demonstrated by measuring the coupling ratio from whole-cell current recordings, independently of the mutant NMDAR membrane expression levels, which were significantly higher for GluN1-GluN2B^V618G^ than GluN1- GluN2B^WT^. In contrast, GluN1-GluN2B^V15M^ showed functional coupling comparable to GluN1-GluN2B^WT^ in spite of its significantly lower membrane abundance. This study constitutes the first detailed characterisation of this V15M mutation and suggests that its pathophysiological impact could be easily explained by low protein abundance and, consequently, reduced membrane expression. This could in turn lead to a decreased formation of NMDAR-BK nanodomains in neurons, although this remains to be explored.

Mutation V618G, however, poses an interesting conundrum. How can a mutation inside the NMDAR pore contribute to the disruption of the functional coupling between GluN1- GluN2B^V618G^ and BK channels? Some pore mutations can destabilise channel openings by altering the receptor conduction pathway (Tristani-Firouzi et al., 2002), while other pore mutants may reshape the pore cavity and alter the channel’s selectivity filter (Cordero-Morales et al., 2006). This may be the case for mutant V618G, with some studies reporting altered Ca^2+^ and Mg^2+^ permeability (Lemke et al., 2014; Vyklicky et al., 2018). However, our results demonstrated that the disruption in NMDAR-BK functional coupling of GluN1-GluN2B^V618G^ could not be ascribed to differences in the permeation of Ca^2+^, as shown with simultaneous Ca^2+^ and voltage-clamp recordings. This is in agreement with previous reports showing comparable Ca^2+^ permeation properties between GluN1-GluN2B^V618G^ and GluN1-GluN2B^WT^ (Fedele et al, 2018). Additionally, we demonstrated that, even if the V618G mutant showed altered Mg^2+^ permeation or block, this could not account for the altered coupling of BK and NMDAR in the nanodomains.

Altogether, our results support the idea that the mechanism underlying the disrupted NMDAR-BK coupling due to mutation V618G is due to a defect of the cell biology of the complex formation. The fact that NMDAR-BK complexes do not form correctly even in the presence of enhanced plasma membrane expression of GluN1-GluN2B^V618G^ reinforces this hypothesis. The combination of PLA and superresolution microscopy demonstrated that complex formation still occurs, albeit with altered size and channel proportions within the nanodomain. This leads us to propose that efficient NMDAR-BK functional coupling requires an adequate proportion between both channels and likely a minimum number of participating units in the nanodomain. Currently, there is very limited information regarding molecular determinants of NMDAR-BK nanodomain formation. Zhang et al. (2018) showed that the S0-S1 loop in the α subunit of BK interacts with intracellular regions of the GluN1 subunit. Our results do not contradict this model, but suggest that GluN2B subunits may also participate in regulating the interaction of NMDAR with BK channels. The question remains how a pore mutation such as V618G may alter such interactions. It is tempting to speculate that this mutation may allosterically disrupt a protein-protein interface that either interacts directly with the BK α subunit or, alternatively, induces changes in GluN1, which in turn alters the interaction with BK. A broader implication of our results is that the presence of different GluN2 regulatory subunits may introduce diversity in the biophysical properties of the nanodomains, and thus in their physiological roles, such as the fine-tuning of synaptic plasticity (Gomez et al., 2021; Zhang et al., 2018). Clearly, a deeper understanding of structural and dynamic properties of NMDAR-BK complex formation is warranted, both from a perspective of their physiological role and as a basis for the pathophysiological consequences of disease-causing mutations.

In summary, we have uncovered a mutation that selectively alters BK-NMDAR complex formation and functional coupling, an effect that may underlie at least some of its pathogenic effects on EIEE27 patients and suggests mechanisms by which BK-NMDAR complexes may modulate synaptic transmission and neuronal function.

