## Supplementary Figures and additional information for "GRIN2B disease-associated mutations disrupt the function of BK channels and NMDA receptor signalling nanodomains"

1. Departamento de Ciencias Medicas Basicas-Fisiologia, Universidad de La Laguna, Tenerife, Spain.
2. Instituto de Tecnologías Biomédicas, Universidad de La Laguna, Tenerife, Spain.
3. Departamento de Bioquímica y Biología Molecular, Universidad de La Laguna, Tenerife, Spain.

SUPLEMENTARY FIGURE 1

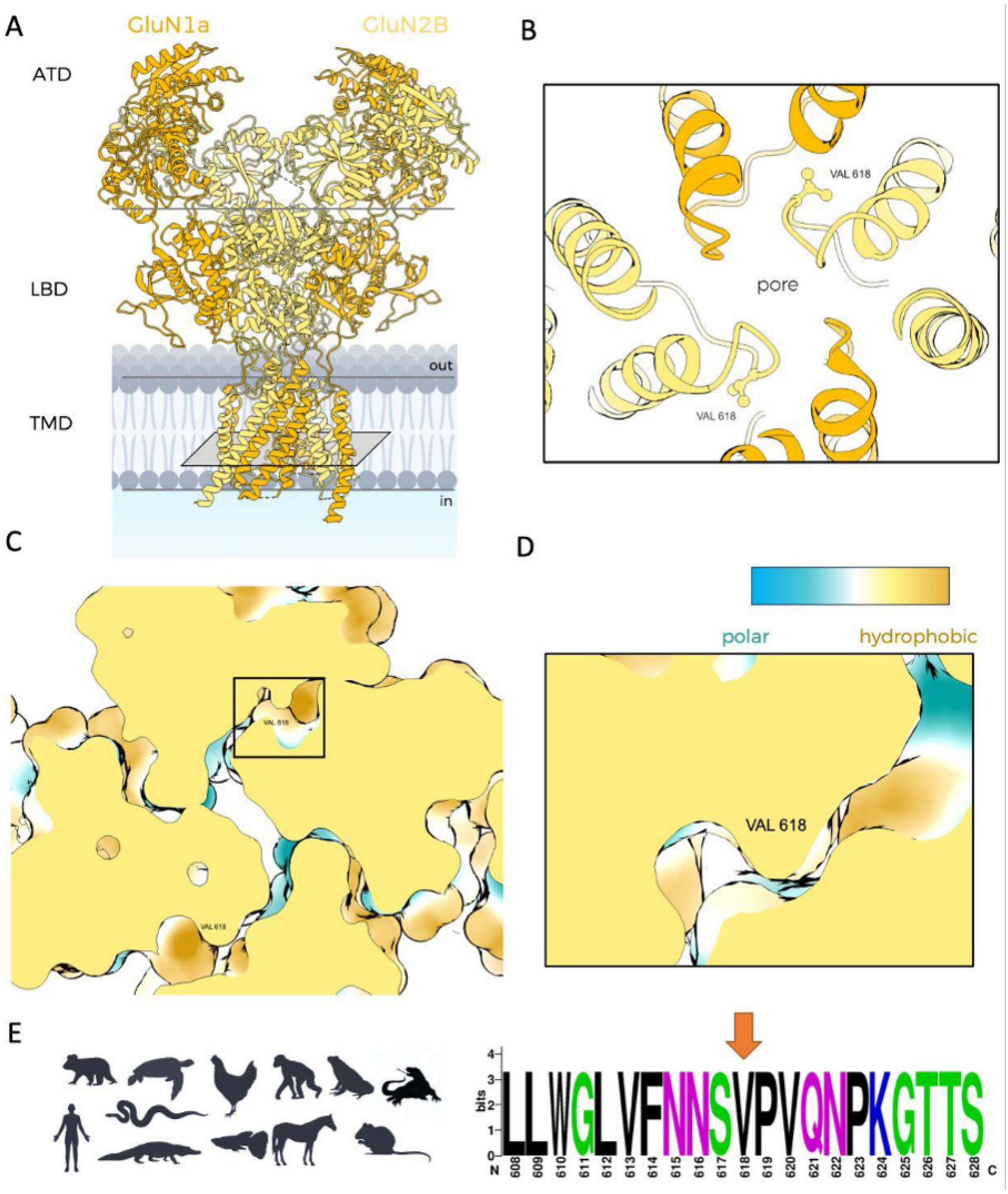


**Supplementary Figure 1**. **Conservation analysis of valine in position 618 in the GluN2B subunit.** (A) Structure of NMDAR composed by GluN1 and GluN2B (pdb: 7SAA). (B) Zoom in on the axial view of the location of valine 618. (C) Hydrophobicity of the environment surrounding V618. Axial view. (D) Zoom in on the hydrophobic profile of V618. (E) V618 and the regions flanking this residue are highly conserved amongst 30 different species from different animal kingdoms, some shown on the left. Amino acids are colored according to their chemical properties: polar amino acids (G, S, T, Y, C, Q, N) are shown in green, basic (K, R, H) in blue, acidic (D, E) in red, and hydrophobic (A, V, L, I, P, W, F, M) amino acids in black. V618 is indicated with an orange arrow. Created with WebLogo plot (University of Berkeley).

SUPPLEMENTARY FIGURE 2

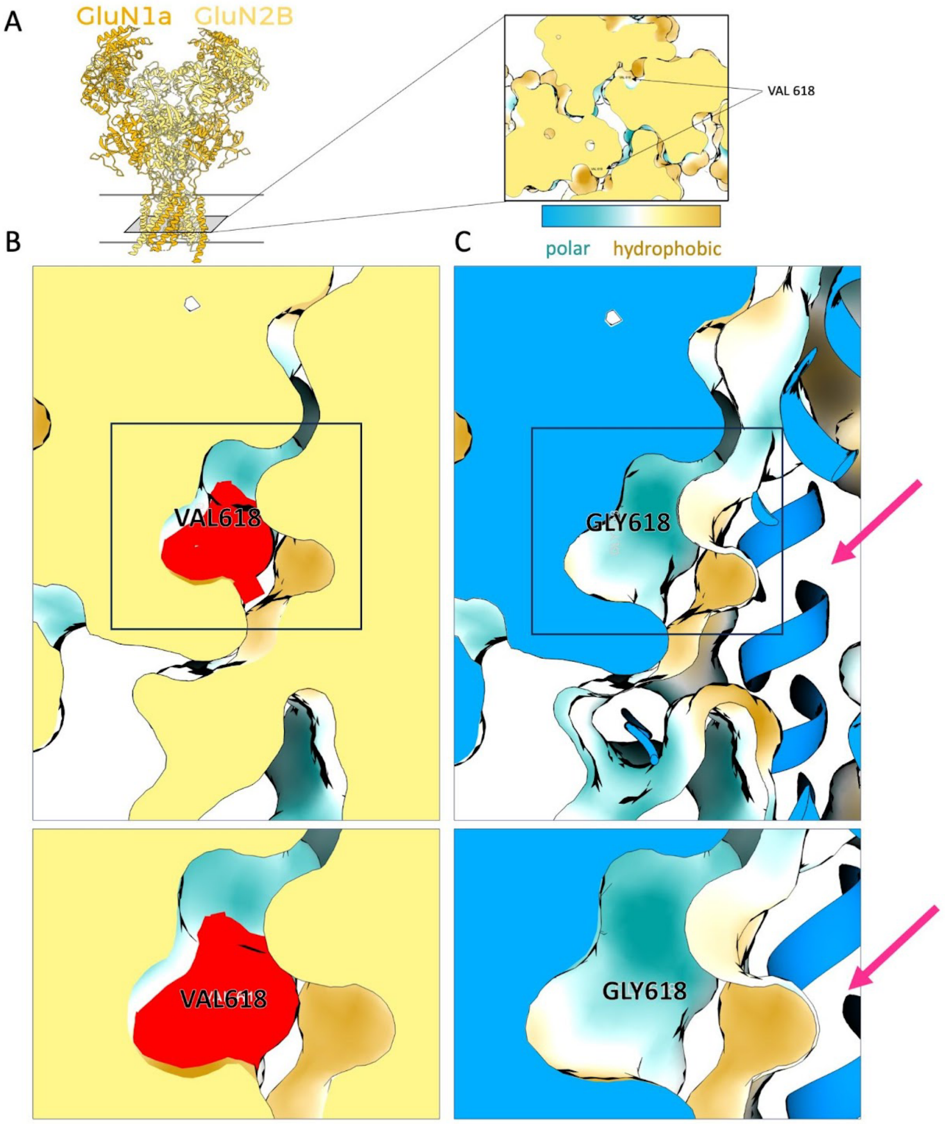


**Supplementary Figure 2. Analysis of the hydrophobicity changes produced by V618G on GluN2B**. (A) Axial view of the location of V618G. (B) Hydrophobicity analysis of V618 revealed the environment in GluN2B$^{WT}$ is highly hydrophobic. (C) Hydrophobicity analysis of mutation V618G showed that the environment surrounding the mutation turned to polar and the creation of a hydrophilic pocket indicated by a pink arrow.

SUPPLEMENTARY FIGURE 3 (WITH ADDITIONAL INFORMATION)

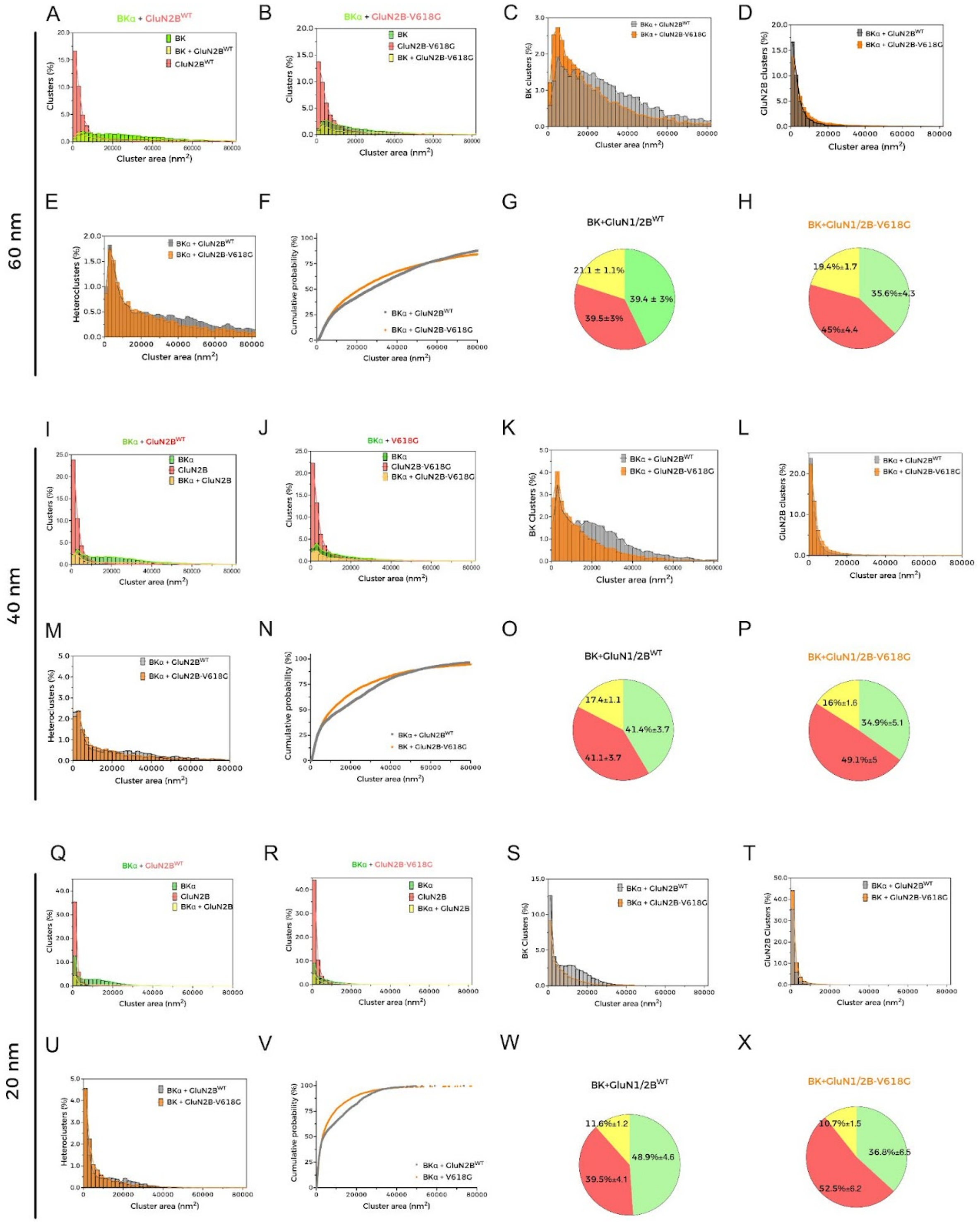


**Supplementary Figure 3. Cluster analysis of BK and GluN2B.** (A) Histograms representing the distribution of clusters in HEK293T cells co-expressing BK and GluN1/2B$^{WT}$ containing BKα alone (green bars), GluN2B$^{WT}$ alone (red bars) or and both proteins (yellow bars). Colored curves outline the histograms to facilitate visualization. Radius cutoff set to 60 nm (r=60nm). (B) Histograms representing the distribution of clusters in HEK293T cells co-expressing BK and GluN1/2B-V618G containing BKα alone (green bars), GluN2B-V618G alone (red bars) or and both proteins (yellow bars), r=60 nm. (C) Comparison of BK homocluster distribution between HEK293T cells co-expressing either BK and GluN1/2B$^{WT}$ (gray bars) or BK and GluN1/2B-V618G (orange bars), r=60nm. (D) Comparison of GluN2B homocluster distribution between HEK293T cells co-expressing either BK and GluN1/2B$^{WT}$ (gray bars) or BK and GluN1/2B-V618G (orange bars), r=60nm. (E) Comparison of heterocluster distribution between HEK293T cells co-expressing either BK and GluN1/2B$^{WT}$ (gray bars) or BK and GluN1/2B-V618G (orange bars), r=60nm. (F) Cumulative probability analysis of heterocluster distribution in cells co-expressing BK and either GluN2B$^{WT}$ or GluN2B-V618G, r=60 nm. (G-H) Pie charts of the % of cluster distribution of individual experiments in cells co-expressing BK and either GluN1/2B$^{WT}$ or GluN1/2B-V618G, r=60 nm. (I) Histograms representing the distribution of clusters in HEK293T cells co-expressing BK and GluN1/2B$^{WT}$ containing BKα alone (green bars), GluN2B$^{WT}$ alone (red bars) or and both proteins (yellow bars). Colored curves outline the histograms to facilitate visualization. Radius cutoff set to 40 nm (r=40nm). (J) Histograms representing the distribution of clusters in HEK293T cells co-expressing BK and GluN1/2B-V618G containing BKα alone (green bars), GluN2B-V618G alone (red bars) or and both proteins (yellow bars), r=40 nm. (K) Comparison of BK homocluster distribution between HEK293T cells co-expressing either BK and GluN1/2B$^{WT}$ (gray bars) or BK and GluN1/2B-V618G (orange bars), r=40nm. (L) Comparison of GluN2B homocluster distribution between HEK293T cells co-expressing either BK and GluN1/2B$^{WT}$ (gray bars) or BK and GluN1/2B-V618G (orange bars), r=40nm. (M) Comparison of heterocluster distribution between HEK293T cells co-expressing either BK and GluN1/2B$^{WT}$ (gray bars) or BK and GluN1/2B-V618G (orange bars), r=40nm. (N) Cumulative probability analysis of heterocluster distribution in cells co-expressing BK and either GluN2B$^{WT}$ or GluN2B-V618G, r=40 nm. (O-P) Pie charts of the % of cluster distribution of individual experiments in cells co-expressing BK and either GluN1/2B$^{WT}$ or GluN1/2B-V618G, r=40 nm. (Q) Histograms representing the distribution of clusters in HEK293T cells co-expressing BK and GluN1/2B$^{WT}$ containing BKα alone (green bars), GluN2B$^{WT}$ alone (red bars) or and both proteins (yellow bars). Colored curves outline the histograms to facilitate visualization. Radius cutoff set to 20 nm (r=20nm). (R) Histograms representing the distribution of clusters in HEK293T cells co-expressing BK and GluN1/2B-V618G containing BKα alone (green bars), GluN2B-V618G alone (red bars) or and both proteins (yellow bars), r=20 nm. (S) Comparison of BK homocluster distribution between HEK293T cells co-expressing either BK and GluN1/2B$^{WT}$ (gray bars) or BK and GluN1/2B-V618G (orange bars), r=20nm. (T) Comparison of GluN2B homocluster distribution between HEK293T cells co-expressing either BK and GluN1/2B$^{WT}$ (gray bars) or BK and GluN1/2B-V618G (orange bars), r=20nm. (U) Comparison of heterocluster distribution between HEK293T cells co-expressing either BK and GluN1/2B$^{WT}$ (gray bars) or BK and GluN1/2B-V618G (orange bars), r=20nm. (V) Cumulative probability analysis of heterocluster distribution in cells co-expressing BK and either GluN2B$^{WT}$ or GluN2B-V618G, r=20 nm. (W-X) Pie charts of the % of cluster distribution of individual experiments in cells co-expressing BK and either GluN1/2BWT or GluN1/2B-V618G, r=20 nm.
